# Evolutionary shortcuts via multi-nucleotide substitutions and their impact on natural selection analyses

**DOI:** 10.1101/2022.12.02.518889

**Authors:** Alexander G Lucaci, Jordan D Zehr, David Enard, Joseph W. Thornton, Sergei L. Kosakovsky Pond

**Affiliations:** Institute for Genomics and Evolutionary Medicine, Temple University, Philadelphia, PA, USA; Dpt. of Ecology and Evolutionary Biology, University of Arizona; Depts. of Human Genetics and Ecology & Evolution, University of Chicago

**Keywords:** Molecular Evolution, Evolutionary shortcuts, Multi-nucleotide substitutions, codon substitution models

## Abstract

Inference and interpretation of evolutionary processes, in particular of the types and targets of natural selection affecting coding sequences, are critically influenced by the assumptions built into statistical models and tests. If certain aspects of the substitution process (even when they are not of direct interest) are presumed absent or are modeled with too crude of a simplification, estimates of key model parameters can become biased, often systematically, and lead to poor statistical performance. Previous work established that failing to accommodate multi-nucleotide (or multi-hit, MH) substitutions strongly biases dN/dS-based inference towards false positive inferences of diversifying episodic selection, as does failing to model variation in the rate of synonymous substitution (SRV) among sites. Here we develop an integrated analytical framework and software tools to simultaneously incorporate these sources of evolutionary complexity into selection analyses. We found that both MH and SRV are ubiquitous in empirical alignments, and incorporating them has a strong effect on whether or not positive selection is detected, (1.4-fold reduction) and on the distributions of inferred evolutionary rates. With simulation studies, we show that this effect is not attributable to reduced statistical power caused by using a more complex model. After a detailed examination of 21 benchmark alignments and a new high-resolution analysis showing which parts of the alignment provide support for positive selection, we show that MH substitutions occurring along shorter branches in the tree explain a significant fraction of discrepant results in selection detection. Our results add to the growing body of literature which examines decades-old modeling assumptions (including MH) and finds them to be problematic for comparative genomic data analysis. Because multi-nucleotide substitutions have a significant impact on natural selection detection even at the level of an entire gene, we recommend that selection analyses of this type consider their inclusion as a matter of routine. To facilitate this procedure, we developed, implemented, and benchmarked a simple and well-performing model testing selection detection framework able to screen an alignment for positive selection with two biologically important confounding processes: site-to-site synonymous rate variation, and multi-nucleotide instantaneous substitutions.

## Introduction

Reliable and robust detection of natural selection from coding sequences continues to be of significant interest in comparative genomics and evolutionary biology literature. Estimation of dN/dS using codon-substitution models (e.g., as implemented in Kosakovsky Pond *et al*. (2020)) is a workhorse of selection detection. Its seminal methods were published several decades ago (Goldman and Yang, 1994; Muse and Gaut, 1994) and still enjoy widespread use. Because computers have gotten faster and datasets much larger, some of the simplifying assumptions in the seminal dN/dS selection tests have been re-examined and generally found to be wanting. For example, the assumption that synonymous rates (dS) do not vary across sites in a gene is nearly universally violated, and it has been shown to inflate the rates of false positive inferences in selection tests (Davydov *et al*., 2019; Pond and Muse, 2005; Wisotsky *et al*., 2020). This finding and its repeated corroborations have led us to recently recommend that synonymous rate variation (SRV) be incorporated into all selection tests as a matter of routine (Wisotsky *et al*., 2020).

Another important modeling assumption is that codon substitutions involving multiple nucleotides (e.g., *ACC→AGG*) must be the result of several evolutionary steps, each of which replaces a single nucleotide (e.g., e.g., *ACC→ACG→AGG*). This assumption is implemented in classic (and still used in the vast majority of applications) tests of selection by setting the instantaneous rates for all multi-nucleotide substitutions to zero. Numerous genetic and molecular studies (Arana *et al*., 2008; Assaf *et al*., 2017; Besenbacher et al., 2016; Chen et al., 2014, 2015; Harris and Nielsen, 2014; Hodgkinson and Eyre-Walker, 2011; Loeb and Monnat, 2008; Matsuda et al., 2000; Prendergast et al., 2019; Saribasak et al., 2012; Schrider et al., 2011; Seplyarskiy et al., 2015; Stone et al., 2012) have established that replication errors often occur at adjacent sites, producing multinucleotide (or multi-hit, MH) mutations that account for ~ 1% of all mutations through several mechanisms, including polymerase *ζ*. Multiple studies have shown that realistic levels of MH mutations can produce uncontrolled rates of false positive inferences of episodic diversifying selection. This is because when a standard model “crams” multiple non-synonymous single-nucleotide substitutions onto a short branch to explain a non-synonymous MH substitution, the *dN/dS* ratio estimate will be artificially inflated to accommodate this. Further, in two genome-wide datasets, MH substitutions within the same codon have been found to account for virtually all the support for episodic diversifying selection, and re-analyzing the data with a model that incorporates MHs as single substitution events dramatically reduces the number of positive inferences of diversifying selection (Venkat *et al*., 2018). A subsequent study confirmed the high rate of false-positive inferences that can be caused by MHs (Dunn *et al*., 2019); another elaborated the MH model and showed that it provides a significantly improved statistical fit to most genes in a collection of > 42,000 empirical alignments (Lucaci *et al*., 2021).

The combined effect of SRV and MH on selection tests has not been systematically investigated, however. Further, no comprehensive approach has yet been developed that simultaneously incorporates both phenomena for testing hypotheses of positive selection. Here we present such a framework: we modified the BUSTED method for episodic diversifying selection (EDS) detection (Murrell *et al*., 2015) to allow synonymous rate variation among codons (+S) and multi-nucleotide substitutions (+MH, including both double- and triple-nucleotide substitutions), and developed model-selection and model-averaging approaches that allow either or both forms of heterogeneity to be incorporated according to their fit to the sequence data. We used this method to re-evaluate the evidence for positive selection in 21 diverse benchmark alignments that historically have played an important role in the development of tests for detecting selection. To understand the power and accuracy of this approach, we applied our framework to both simulated data and a large alignment of 9,181 of proteins in orthologs of 24 mammlian species, previously studied by Enard *et al*. (2016) in the context selection as a result of interactions with viral proteins. Our practical framework can be used to balance the impetus to prevent false positive inferences of selection when S and/or MH effects are present but not incorporated, and the desire to prevent the loss of statistical power, which can occur when models of greater parametric complexity are used.

## Results

### High-level model description

We compared four different BUSTED class models (see Methods for complete details), each of which tests for evidence of a non-zero fraction of branch-site combinations evolving with *ω*> 1, but makes different assumptions about confounding evolutionary processes. We evaluate four models:

1. the baseline model (BUSTED),
2. a model which adds site-to-site synonymous rate variation (+S),
3. a model with support for instantaneous double- and triple-nucleotide substitutions within a single codon (+MH),
4. and a model with support for both (+S+MH),

(cf Table 1 for additional details). These models form a nested hierarchies (*BUSTED*⊂+*S*⊂+*S*+*MH* and *BUSTED*⊂+*MH*⊂+*S*+*MH*) and can be compared using either information theoretic criteria or pairwise likelihood ratio tests.

**Table 1.** Substitution models considered in this paper. *B* - the number of branches in the phylogenetic tree. *K* and *L* are user-tunable parameters, set to 3 each by default.

| Model | Reference | Non-synonymous rates | Synonymous rates | Multi-nucleotide substitutions | Number of parameters |
| --- | --- | --- | --- | --- | --- |
| BUSTED | Murrell *et al.* (2015) | Random effects branch-site modeled by a K(=3)-bin discrete distribution | None | None | $B + 13 + 2 \times K$ |
| +S | Wisotsky *et al.* (2020) | Random branch-site effects modeled by a K(=3)-bin general discrete distribution | Random site effects modeled by an L(=3)-bin unit mean general discrete distribution | None | $B + 11 + 2 \times (K + L)$ |
| +MH | Lucaci *et al.* (2021) | Random branch-site effects modeled by a K(=3)-bin general discrete distribution | None | Alignment-wide double- ($\delta$) and triple- ($\psi$) nucleotide substitution rates | $B + 15 + 2 \times K$ |
| +S+MH | This paper | Random branch-site effects modeled by a K(=3)-bin general discrete distribution | Random site effects modeled by an L(=3)-bin unit mean general discrete distribution | Alignment-wide double- ($\delta$) and triple- ($\psi$) nucleotide substitution rates | $B + 13 + 2 \times (K + L)$ |

### Analysis of benchmark alignments

It is informative to begin by examining how the four competing models (Table 2) perform on a collection of empirical sequence alignments. We screened 21 alignments for evidence of EDS. These alignments were chosen because they have each been previously analyzed (many in multiple papers) for evidence of natural selection using a variety of models, and because they represent different alignment sizes, diversity levels, and taxonomic groups, all of which impact selection analyses.

**Table 2.**
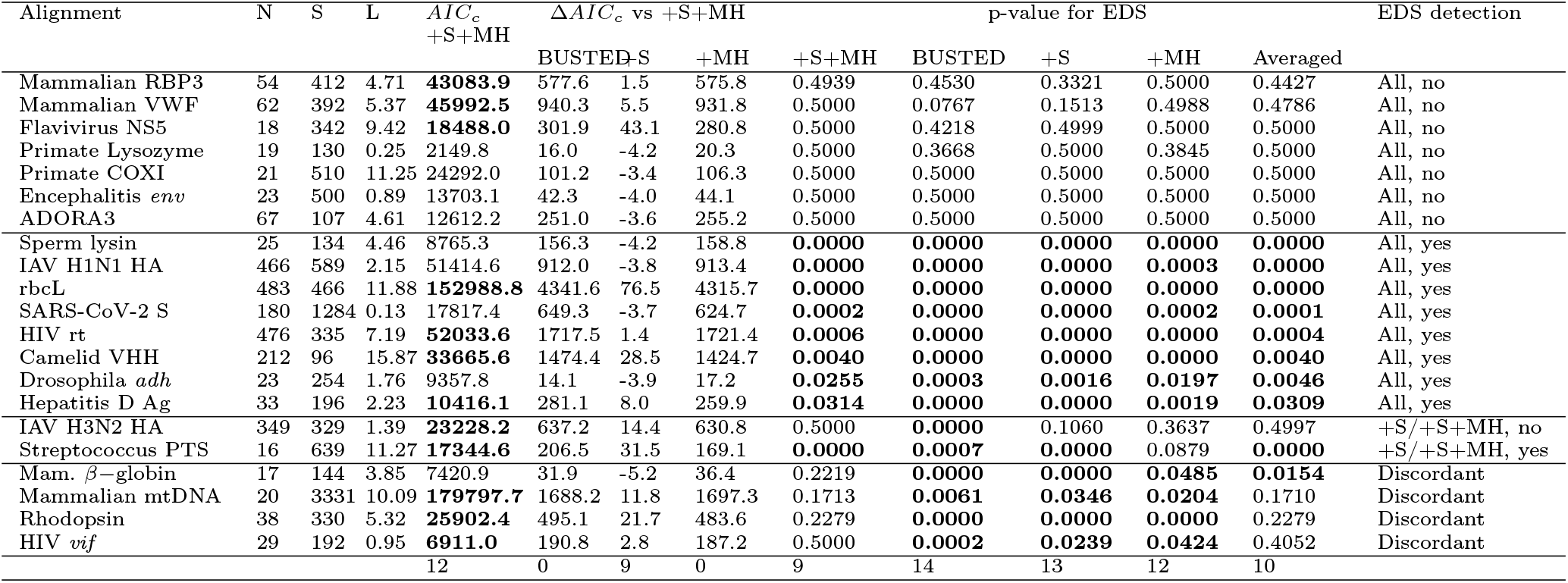
Selection analysis on benchmark alignments (arranged by EDS detection class first and further sorted by model-averaged p-value). *N* - the number of sequences, *S* - the number of codons, *L* - total tree length in expected substitutions/nucleotide, measured under the BUSTED+S+MH model. *AIC_c_* S+MH - small sample AIC score for the BUSTED+S+MH model (shown in boldface if this model is the best fit for the data, i.e. has the lowest *AIC_c_* score), Δ*AIC_c_* - differences between the *AIC_c_* score for the corresponding model and the BUSTED+S+MH. p-value for ESD: the likelihood ratio test p-value for episodic diversifying selection under the corresponding model (4 digits of precision); shown in boldface if ≤0.05. The Averaged column shows model averaged p-values (see text). The last column indicates model agreement with respect to detecting ESD at *p*≤0.05. The last row shows the number of times each model was preferred by *AIC_c_*, and the number of significant LRT tests for each model and the model averaged approach.

The inclusion of site-to-site synonymous rate variation is strongly supported for all 21 datasets (in agreement with Wisotsky *et al*. (2020)), and further addition of multi-nucleotide substitution (MNS) support is preferred by *AIC_c_* in 12/21 datasets (Table 2). The addition of model complexity reduces the rate at which EDS is detected, with the simplest model (BUSTED) returning significant test results for 14/21 datasets, and the most complex model (+S+MH) – for 9/21. Because our primary analytical endpoint is the detection of EDS, we can categorize the alignments into those where models agree, and those where they disagree. We begin with the **seven** datasets where all four models arrived at a negative result. Some of these examples confirm previous negative findings, and some contradict previous positive (potentially weak) findings.

#### Primate lysozyme

##### (best model: +S)

A version of this dataset was originally used to show lineage specific variation in dN/dS (or *ω*) in Yang (1998), where tests assuming no site-to-site rate variation (SRV) also identified positive selection (mean *ω* > 1) on the hominoid lineage. This evidence is no longer statistically significant if a suitable multiple testing correction is applied to the original results. Overall, this is a low divergence dataset with relatively few substitutions (see Tables 2 and 3).

**Table 3.** Substitution process characterization on benchmark alignments. *ω*3(*p*3) - the maximum likelihood estimate of the *ω* ratio for the positively selected class, along with its estimated fraction, for +S and +S+MH models. *CV*(*α*) - coefficient of variation for the inferred distribution of site-to-site synonymous substitutions rates (+S+MH model). *δ* – the MLE for the two-hit substitution rate, *ψ* – the MLE for the three-hit substitution rate; 0 point estimates are shown as - for readability. Substitutions - the counts (and fractions of total branch × sites pairs) where one (1H), two (2H) or three (3H) nucleotides change along the branch under the +S+MH model. *L* - mean branch lengths for branches experiencing 1H, 2H, and 3H substitutions under the +S+MH model.

| Alignment | $\omega_3(p_3)$ | | $CV(\alpha)$ | $\delta$ | $\psi$ | Substitutions (%) | | | L | | |
| --- | --- | --- | --- | --- | --- | --- | --- | --- | --- | --- | --- |
|  | +MH | +S |  |  |  | 1H | 2H | 3H | 1H | 2H | 3H |
| β-globin | 2.834 (6.08%) | 8.925 (3.70%) | 1.263 | 0.241 | - | 526 (11.78) | 110 (2.46) | 24 (0.54) | 0.26 | 0.36 | 0.52 |
| Primate Lysozyme | 1.002 (0.00%) | 1.002 (0.00%) | 1.242 | - | - | 81 (2.08) | 3 (0.08) | 0 (0.00) | 0.02 | 0.04 | 0.00 |
| Sperm Lysin | 17.459 (7.57%) | 17.020 (7.70%) | 0.867 | - | - | 514 (8.16) | 149 (2.37) | 33 (0.52) | 0.18 | 0.22 | 0.30 |
| HIV vif | 1.226 (1.00%) | 2103.279 (0.05%) | 1.049 | 0.004 | 0.163 | 446 (4.30) | 20 (0.19) | 5 (0.05) | 0.03 | 0.04 | 0.04 |
| Drosophila adh | 4.056 (2.38%) | 4.144 (2.50%) | 0.594 | 0.032 | - | 693 (6.34) | 66 (0.60) | 14 (0.13) | 0.10 | 0.15 | 0.16 |
| Flavivirus NS5 | 1.105 (0.00%) | 1.006 (2.23%) | 1.286 | 0.377 | 0.986 | 1956 (17.33) | 270 (2.39) | 58 (0.51) | 0.48 | 0.58 | 0.58 |
| Hepatitis D Ag | 11.306 (1.71%) | 16.249 (1.96%) | 0.902 | 0.143 | - | 665 (5.39) | 122 (0.99) | 14 (0.11) | 0.08 | 0.13 | 0.18 |
| ADORA3 | 1.013 (0.00%) | 1.000 (3.46%) | 0.662 | 0.053 | - | 1135 (8.35) | 75 (0.55) | 3 (0.02) | 0.06 | 0.09 | 0.14 |
| Streptococcus PTS | 9.489 (1.56%) | 11.871 (1.85%) | 1.046 | 0.310 | 1.054 | 1245 (6.72) | 293 (1.58) | 76 (0.41) | 1.79 | 1.98 | 1.96 |
| Mammalian COXI | 1.021 (1.06%) | 1.000 (1.09%) | 2.342 | - | - | 3160 (15.89) | 132 (0.66) | 6 (0.03) | 0.43 | 0.45 | 0.59 |
| Encephalitis env | 3.166 (0.00%) | 1.002 (0.00%) | 0.671 | 0.012 | - | 1068 (4.97) | 26 (0.12) | 0 (0.00) | 0.04 | 0.05 | 0.00 |
| Rhodopsin | 5.453 (0.37%) | 6.376 (1.31%) | 1.403 | 0.345 | 0.515 | 2488 (10.77) | 257 (1.11) | 45 (0.19) | 0.12 | 0.15 | 0.16 |
| Camelid VHH | 9.193 (2.53%) | 24.513 (2.04%) | 0.817 | 0.157 | - | 2393 (6.91) | 528 (1.52) | 72 (0.21) | 0.09 | 0.11 | 0.12 |
| Mammalian RPB3 | 1.258 (0.44%) | 1.541 (2.03%) | 0.593 | 0.111 | 0.054 | 4093 (9.46) | 300 (0.69) | 18 (0.04) | 0.08 | 0.10 | 0.11 |
| Mammalian VWF | 1.073 (0.36%) | 1.973 (2.35%) | 0.643 | 0.130 | - | 4608 (9.71) | 381 (0.80) | 22 (0.05) | 0.08 | 0.11 | 0.17 |
| Mammalian mtDNA | 1.310 (1.04%) | 1.434 (1.33%) | 1.268 | 0.227 | - | 19892 (16.14) | 1873 (1.52) | 225 (0.18) | 0.42 | 0.57 | 0.67 |
| IAV H3N2 HA | 1.002 (0.00%) | 1.550 (29.48%) | 1.064 | 0.062 | 0.015 | 1320 (0.74) | 29 (0.02) | 1 (0.00) | 0.00 | 0.00 | 0.01 |
| HIV rt | 50.714 (0.07%) | 48.230 (0.10%) | 0.940 | 0.028 | - | 4149 (1.35) | 129 (0.04) | 10 (0.00) | 0.02 | 0.02 | 0.03 |
| rbcl | 37.569 (0.14%) | 49.827 (0.19%) | 0.831 | 0.113 | 0.016 | 12185 (2.77) | 653 (0.15) | 72 (0.02) | 0.02 | 0.03 | 0.02 |
| SARS-CoV-2 S | 5.990 (20.12%) | 5.746 (29.04%) | 3.132 | 0.012 | - | 421 (0.13) | 13 (0.00) | 0 (0.00) | 0.00 | 0.00 | 0.00 |
| IAV H1N1 HA | 1039.054 (0.01%) | 862.821 (0.01%) | 0.835 | - | - | 3216 (0.73) | 43 (0.01) | 2 (0.00) | 0.01 | 0.02 | 0.00 |

#### Tick-borne flavivirus NS-5 gene

##### (+S+MH)

This dataset was analyzed in Yang *et al*. (2000a) and originally sourced from Kuno *et al*. (1998); no evidence of positive selection was found in the original papers. This is a high-divergence alignment, including 51 events when all three nucleotides are inferred to have changed along a single branch at a particular site (Table 3).

#### ADORA3

##### (+S)

This alignment of adenosin A3 receptor (placental mammals) was analyzed using Bayesian mutation selection models by Rodrigue *et al*. (2021), who reported weak to no evidence of adaptive evolution.

#### COXI

##### (+S)

Primate cytochrome oxidase subunit I mitochondrial sequences were previously analyzed in (Seo *et al*., 2004) using Bayesian methods; they found significant lineage-to-lineage variation in absolute synonymous and non-synonymous rates, but strong conservation (ω ≪ 1) overall. We find no evidence of MNS, including 0 point estimates for *δ* and *ψ*, despite > 100 events with more than one nucleotide being substituted along a single branch at a given site (Table 3). As reported by (Lucaci *et al*., 2021), standard models are often able to properly account for multiple nucleotide substitution events along long branches.

#### Japanese encephalitis env gene

##### +S)

This alignment was included in Yang *et al*. (2000a), who found it to be subject to strong purifying selection.

#### VWF

##### (+S+MH)

The von Willbrand factor gene (placental mammals) from Rodrigue *et al*. (2021), who found some evidence of positive selection with mutation-selection Bayesian models (none with standard site-heterogenous codon models), although the authors caution that other un-modeled evolutionary processes (e.g. CpG hyper-mutability) could confound inference.

#### RBP3

##### (+S+MH)

Retinol-binding protein 3 (placental mammals) from Rodrigue *et al*. (2021); no evidence of positive selection was found in this gene by the original authors.

Next, we describe the **eight** datasets where all of our models found statistical evidence for EDS (LRT *p* ≤ 0.05).

#### adh

##### (+S)

Drosophila alcohol dehydrogenase (adh) gene (originally from Hudson *et al*. (1987)), studied in numerous selection detection papers, including Yang *et al*. (2000a) and Rodrigue *et al*. (2021). Most analyses failed to detect evidence of diversifying selection, despite long-standing supposition that balancing selection is acting on this gene. Rodrigue *et al*. (2021) reported that mutation-selection models detected numerous sites subject to selection; our methods allocate 2.5% (of branch-site pairs) to the positively selected regime.

#### Lysin

##### (+S)

This alignment of abalone sperm lysin from Yang *et al*. (2000b) is a canonical example of diversifying positive selection, e.g., due to self-incompatibility constraints. There is no support for MNS in this alignment despite relatively high divergence and numerous multi-nucleotide branch-site substitution events (Table 3).

#### Hepatitis D Ag

##### (+S+MH)

Anisimova and Yang (2004) analyzed an alignment of Hepatitis Delta virus antigen gene with site-heterogeneous methods, and reported extensive positive selection. Our best fitting model (+S+MH) estimates 1.7% fraction of branch-site pairs to be subject to EDS (*ω*≈ 11.3). The MNS signal is entirely due to double-nucleotide substitutions 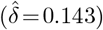. While all models have *p*≤0.05 for EDS, the p-value is highest for the +S+MH model, as we show later, this is a common pattern, when the addition of MNS support reduces (or eliminates) statistical significance levels of tests for EDS.

#### Camelid VHH

##### (+S+MH)

Su *et al*. (2002) studied this variable regions of immunoglobulin heavy chains in camelids using relatively underpowered counting methods, and found extensive evidence of positive selection. The best fitting model (+S+MH) allocates 2.5% of branch-site pairs to the positively selected class (*ω* ≈ 9.2) and the MNS signal is driven by double-nucleotide substitutions 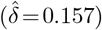.

#### HIV-1 rt

##### (+S+MH)

A HIV-1 reverse transcriptase alignment comprises pairs of sequences from individuals prior to and following antiretroviral treatment, studied by Seoighe *et al*. (2007) to examine selective pressures due to the development of drug resistance. There is marginal evidence of MNS based on *AIC_c_*, and very strong (*ω* ~ 50) positive selection on a small (~ 0.1%) fraction of branch-site pairs.

#### rbcL

##### (+S+MH)

Tamuri and Dos Reis (2022) examined this alignment of plant RuBisCO with a penalized likelihood mutation-selection model (no MNS), and identified numerous sites subject to pervasive positive selection. We find strong evidence of MNS based involving both two- and three-nucleotide substitutions, and very strong (*ω* ~ 38) positive selection on a small (~ 0.14%) fraction of branch-site pairs.

#### SARS-CoV-2 S

##### (+S)

A collection of full-length SARS-CoV-2 spike genes from variants of concern and other representative lineages, obtained from GISAID (Shu and McCauley, 2017). Numerous previous studies (e.g. Martin *et al*. (2021, 2022); Viana *et al*. (2022)) detected positive selection on this gene, driven primarily by immune selective pressure and enhanced transmissibility. The best-fitting model (+S) infers that a very large fraction of this gene (~ 20%) is subject to positive selection (*ω* ≈ 5.7).

#### IAV H1NA1

##### (+S)

Tamuri and Dos Reis (2022) performed a detailed analysis of this H1N1 Influenza A virus (human hosts) hemagluttinin dataset and found 14-18 (depending on model parameters) sites under selection. Our analysis detects very strong (*ω* ~ 1000) positive selection on a very small (~ 0.01%) fraction of branch-site pairs, and no evidence of MNS.

Lastly, we discuss the **six** remaining datasets, where EDS detection depends on the model. These are the most important to address, because they represent the cases where selection inference is, in some sense, not robust to modeling assumptions.

#### *β*—globin

##### (+S)

Mammalian *β*—globin is one of the datasets from Yang *et al*. (2000a) where positive selection has been inferred, and confirmed using many other studies and methods (e.g., Rodrigue *et al*. (2021)). All of our models, except for +S+MH, including the best fitting model (+S), also infer positive selection. However, the addition of MNS (+S+MH) model results in the elimination of statistical significance; this appears to be the case of overfitting, because +S+MH is supported neither by *AIC_c_*, nor by direct nested LRT between the two models (*p*~0.5).

#### HIV-1 vif

##### (+S+MH)

HIV-1 viral infectivity factor (*vif*) was inferred to be under positive selection in Yang *et al*. (2000a), but not according to our best-fitting model (+S+MH). The second best fitting model (+S), whose *AIC_c_* is only slightly higher, returns a significant p-value for EDS. To better understand which features of the dataset leads to discordant conclusions, we applied fit profiling techniques (see Methods), and found that a single codon in the alignment (codon 6) contributes the majority of the cumulative likelihood ratio test signal (Figure S1 for the +S model. Furthermore, a single triple-nucleotide substitution along a terminal tree branch at that site, `CAG (Q) → GCA (A)`, contributes the bulk of statistical support for EDS in the +S model and the addition of MHS the model completely eliminates this support (Figure 1.A). Multi-nucleotide substitutions along short branches have been shown to return false positive selection detection results in simulations (Lucaci *et al*., 2021; Venkat *et al*., 2018). Masking a single codon (GCA) with gaps in the alignment and rerunning BUSTED+S yields a non-significant p-value for EDS. The fact that a single codon can be responsible for the detection of gene-wide positive selection does not inspire confidence in the positive result with the +S model.

**FIG. 1.**
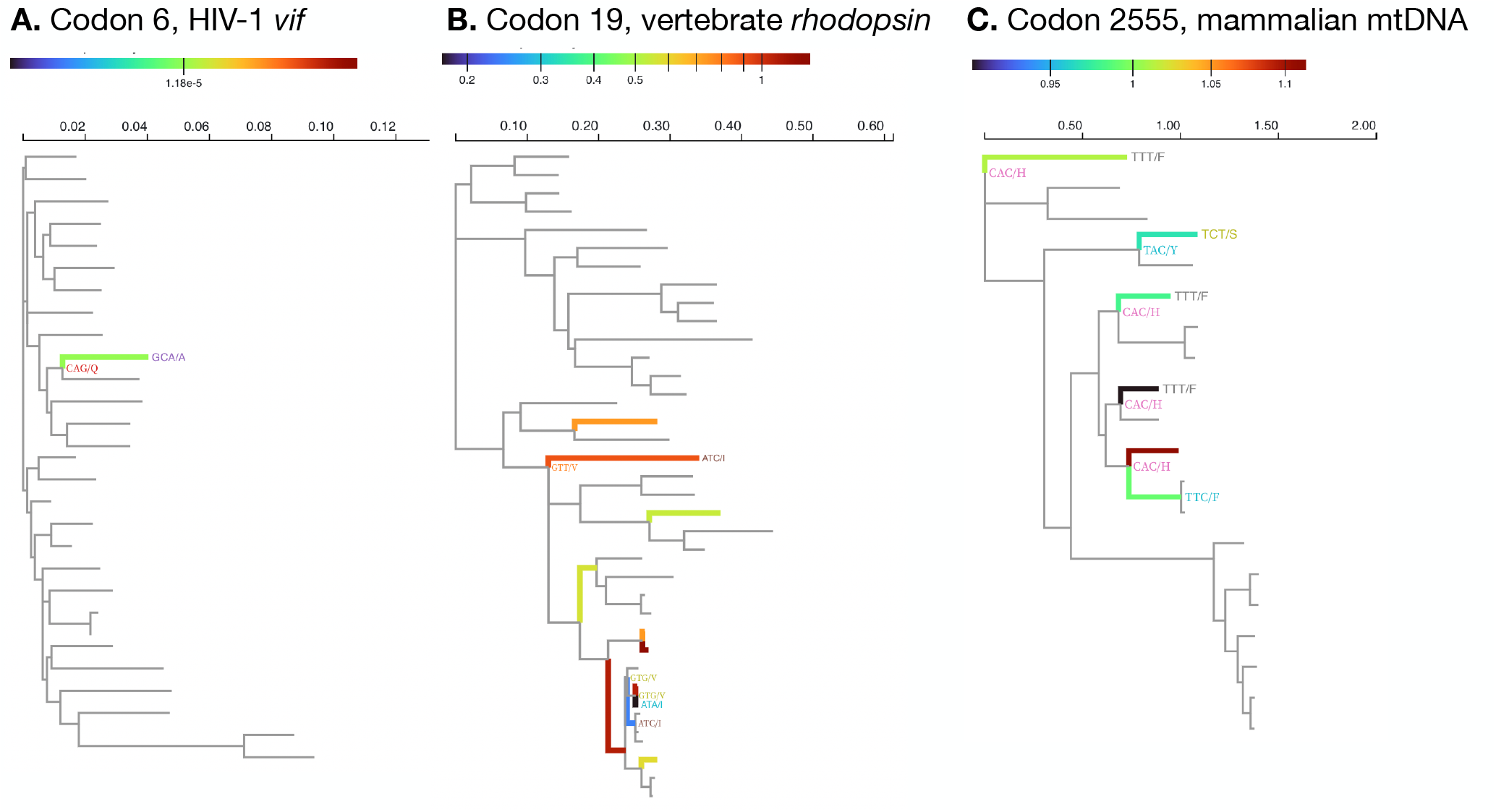
Example sites from benchmark alignments with discordant selection signal. Only substitutions involving multiple nucleotides are labeled (codon/amino-acid). Coloring of the branches represents the ratio of empirical Bayes factors for *ω*> 1 at this branch/site (see Methods) between the +S+MH and +S models. Values < 1 imply that the +S+MH model has less support for *ω* > 1 than the +S model. The scales are different for each of the examples because they have dramatically different ranges.

#### Streptococcus

##### (+S+MH)

Dunn *et al*. (2019) analyzed this trehalose-specific PTS sugar transporter system alignment (gene 2 in their study) using parameter-rich models including MNS and found evidence of positive selection (*ω*_+_ = 4.9,*p*_+_ =0.028). Our best fitting model (+S+MH) infers a 1.6% fraction of branch-site pairs to be subject to EDS (*ω*≈9.5), and so does the second best fitting model (+S). The contrarian model (+MH) is a much poorer fit to the data, and can be discounted.

#### Vertebrate Rhodopsin

##### (+S+MH)

This dim-light vision protein was exhaustively analyzed by Yokoyama *et al*. (2008) with comparative methods and via experimental assays. They found that amino-acid substitutions at 12 sites altered a key phenotype (absorption wavelength, λ_max_ of some sequences, but that traditional site-level methods for diversifying selection detection found fewer sites without significant phenotypic impact. Our best (+S+MH) and second best (+S) fitting models return strongly discordant results for EDS (*p*=0.23 and *p*<0.0001, respectively). Both double- (257 events) and triple-nucleotide (45 events) substitution rates have non-zero MLE (Table 3). Compared to the +S model, the +S+MH model infers a smaller fraction of branch site combinations (0.37% vs 1.31%) with lower *ω* (5.5 vs 6.4). We noticed a similar trend with simpler rate variation models in Lucaci *et al*. (2021) – the inclusion of MNS reduces *ω* estimates. Most of the sites which contribute signal to EDS detection with the +S model, contribute less (or no) signal under the +S+MH model (Figure S1), with strong reduction occurring at short branches which harbor multinucleotide substitutions (Figure 1.B). One obvious explanation for model discordance is loss of power for the more parameter rich +S+MH model, but it seems unlikely. When we simulate under the +S model (using parameter fits from the data, which includes EDS), the power to detect selection is comparable between the models (0.99 for +S+MH vs 1.00 for +S, please see the Simulated Data section for more details).

#### Mammalian mtDNA

##### (+S+MH)

This concatenated alignment of mammalian mitochondrial genomes ships as a test dataset with the PAML package and has been recently re-analyzed by Jones *et al*. (2018) using several models including those supporting MNS, which were preferred. Our analyses also indicate support for MNS (both double- and triple-nucleotide), but the +S+MH model (best-fitting) and +S model (second best fitting) disagree on the presence of EDS. The +S model (Table 3) allocates 1.3% of branch-site combinations to a weakly selected component (*ω* = 1.4), but the source of this support is diffuse across many sites, with relatively little signal contributed by individual sites (Figure S1). Similarly, the reduction is EDS support under +S+MH is also diffuse and less pronounced for individual sites. Because of longer branches, even sites with extensive MNS have a minor decrease in inferred local support for EDS when comparing +S+MH and +S models (Figure 1.C). Analysis of 100 replicates generated under the +S model shows that the lack of detection under S+MH is probably not because of significant power loss (0.50 for +S+MH vs 0.65 for +S, please see the Simulated Data section for more details)

#### IAV H3N2

##### (+S+MH)

Yang (2000) examined an alignment of human isolates of H3N2 Influenza A virus hemagluttinin sequences, originally studied by Bush *et al*. (1999), for evidence of EDS using site-level methods and found support for it. With the exception of the poorly-fitting BUSTED model, our analyses fail to find evidence of EDS, potentially because of extensive synonymous rate variation (Wisotsky *et al*., 2020), although the addition of MNS support without SRV (+MH model), also removes the selection signal.

In summary, there is a good degree of agreement between models in detecting episodic diversifying selection on 21 benchmark datasets, with 15/21 agreements among all models and 17/21 for the best fitting models (+S and +S+MH), Cohen’s *κ* = 0.63 (substantial agreement). In all four substantively discordant cases, +S+MH did not find evidence for selection, while +S – did find such evidence. This greater “conservatism” on the part of +S+MH is unlikely to be due to significant loss of power relative to +S (see Simulations), and manual examination of discordant datasets points towards events which involve multi-nucleotide changes along shorter tree branches as a main driver of the differences.

In nearly all of the datasets, +S+MH model infers a smaller proportion of sites subject to weaker (smaller *ω*) selection, implying that the +S model, at least for the datasets where +S+MH is preferred by *AIC_c_* may be absorbing some of the unmodeled multi-nucleotide substitutions into the *ω* distribution (Jones *et al*., 2018; Lucaci *et al*., 2021).

###### Model averaged p-values

As a simple and interpretable approach to synthesize the results different models fitted to the same dataset, and account for different goodness-of-fit, we propose a model averaged p-value. It is defined as 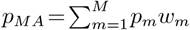, where the sum is taken over all models considered, *p_m_* is the p-value returned by model *m* and *w_m_* is the Akaike weight for model *m* (Wagenmakers and Farrell, 2004). 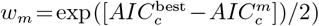, where 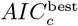 is the score of the best-fitting model normalized to sum to 1 over all *M* models. The Akaike weight, *w_m_*, can be interpreted as ~ *P* (model = m|data), when *M* models are being compared. Consequently, if model *m* returns the likelihood ratio test of *LRT_m_*, then 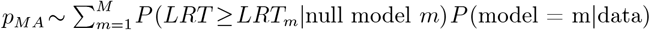.

The model averaged approach detects the same 9 datasets as the +S+MH model, and also the *β*—globin dataset, where the +S model (with EDS signal) has a sufficiently significant edge in goodness of fit to “outvote” the +S+MH model (Table 2). As our simulation results show (next section), the model averaged approach is a simple and automated way to control false positives, while maintaining very good power.

### Analysis of simulated alignments

#### Four taxon tree null simulations

We generated synthetic alignments of 4 sequences with 800 codons each, using the tree shown in Figure 2, subject to negative selection or neutral evolution (*ω*_1_ =0.1(50%), *ω*_2_ = 0.5(25%),*ω*_3_ = 1.0(25%)), under the +S or +S+MH models. We varied the 2H rate (*δ*), and the 3H rate (*ψ*) as well as lengths of 2 of the 5 branches in the tree, generating 100 replicates for each parameter combination considered. We then fitted +S and +S+MH models to all of the replicates, and tabulated false positive rates (FPR).

**FIG. 2.**
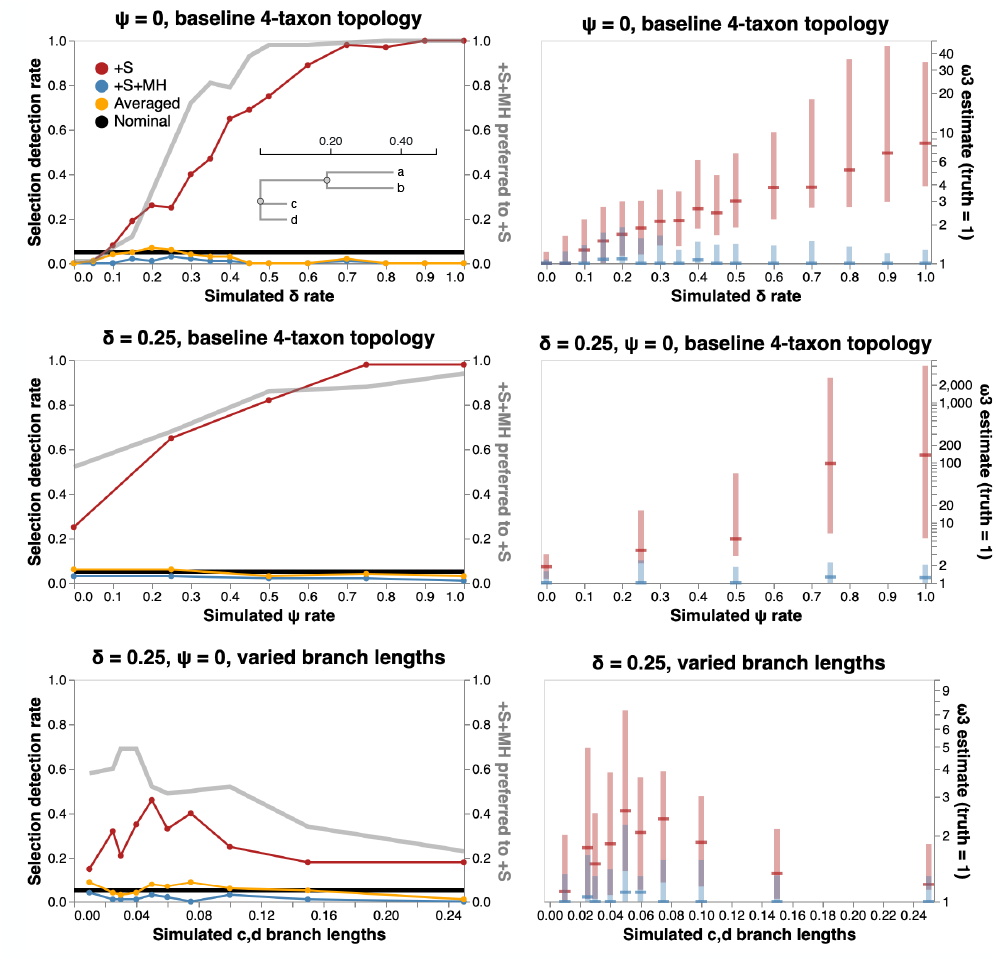
Model performance on null simulated data. Left column: false positive detection rate for EDS (at *p*≤0.05) as a function of rate parameters and branch lengths, and the rate at which +S+MH is preferred to +S by a nested LRT test. Right column: *ω*_3_ estimates (median, IQR) for various simulation scenarios. 100 replicates were generated using the four-taxon tree shown as inset in the top left plot for each parameter combination. For the bottom row, we varies the lengths of branches leading to c and d in the tree

##### False positive rates

As the 2H rate (*δ*) increases (Figure 2), the +S model shows progressively higher FPR (reaching 100%), coupled with increasingly biased estimates of *ω*_3_ – the positive selection model component. On the other hand, the +S+MH model shows nominal or conservative FPR, and generally consistent estimates of *ω*_3_. Because the +S+MH model has increasingly better fit to the data as *δ* becomes larger, the model averaged p-value (which gives progressively more weight to +S+MH), also has controlled FPR, with the exception of slightly elevated rates for 0.15≤*δ*≤0.25. Therefore, the +S model appears to “absorb” unmodeled multiple-hit substitutions into biased *ω* estimates, which leads to catastrophically high rates of false positives. An identical pattern is observed for a fixed *δ* and increasing rates of 3H substitutions (*ψ*), seen in Figure 2. Finally, FPR of the +S model also depends on branch lengths of the tree. In these simple simulations branch lengths ~ 0.05 show an elevation in +S FPR rates. The intuition is simple: very short branches do not accumulate many substitutions (no signal), sufficiently long branches do not benefit as much from access to instantaneous 2H substitutions, because over longer branches it is nearly as easy to obtain a 2H substitution via two (or more) consecutive 1H substitutions allowed in the standard models. Short branches with multi-nucleotide substitutions force the +S model to absorb these unmodeled changes into the *ω* rate, and have the largest effect on FPR rates.

##### Power

On the same 4-taxon tree, we next simulated alignments with a non-zero fraction of the alignment subject to EDS, with the distribution of rates (*ω*_1_ = 0.1(50%),*ω*_2_ = 0.5(40%),*ω*_3_ > 1(10%). We iterated *ω*_3_ over the set {1.25,1.5,2,4,8,16}, set *ψ* = 0, and iterated *δ* over the set {0,0.1,0.2,0.3,0.4,0.5}, for a total of 36 simulation scenarios. The three methods for EDS detection (+S, +S+MH, and model averaged), all gained power as the effect size (*ω*_3_) increased, reaching 100% (Figure 3). When no multi-plehits are allowed (*δ* = 0), +S+MH shows a small loss of power compared to the +S model, but because +S has better fit in nearly every dataset, model averaging rescues most of the power. Both +S and +S+MH return consistent estimates of *ω*_3_. For *δ*> 0 and for *ω*_3_ < 8 the +S model has progressively higher power, but that power comes at the cost of progressively more and more biased estimates of *ω*_3_. This behavior mirrors what we saw for null data, except, for data simulated with positive selection (low or moderate effect sizes), the bias results in a desirable outcome (higher power). Model averaging becomes less effective as δ grows, because the +S model loses goodness-of-fit compared to the +S+MH model. Increasing the fraction of alignments subject to selection to 25% (*ω*_1_=0.1(50%),*ω*_2_=0.5(25%),*ω*_3_>1(25%)) shows the same qualitative behavior, except that all methods have higher power for a given value of *δ* and *ω*_3_ (Figure 3, Figure S2).

**FIG. 3.**
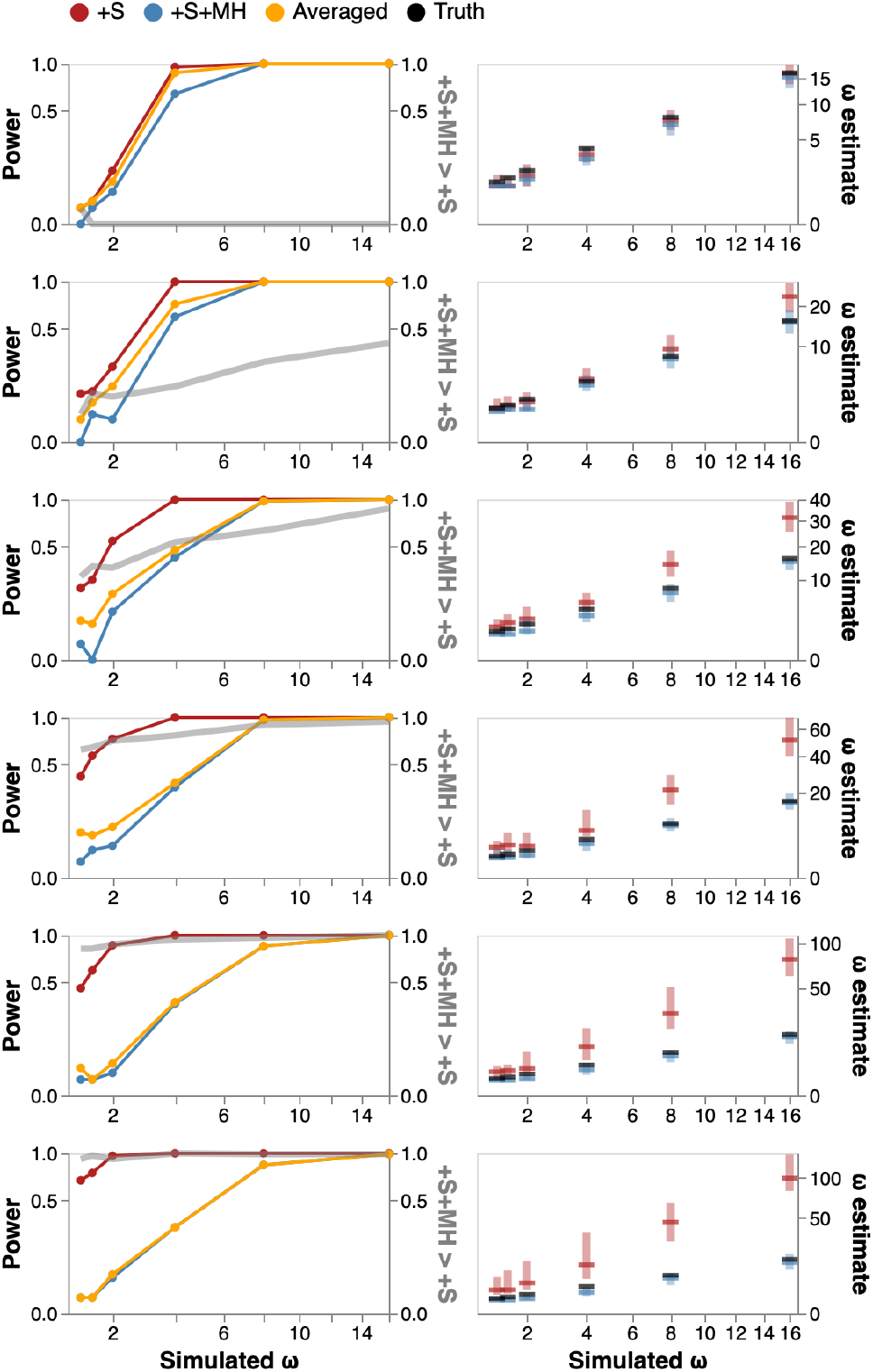
Model performance on data simulated with EDS. Left column: detection rate for EDS (at *p*≤0.05) as a function of rate *ω*_3_ (effect size) and *δ* (confounding parameter), and the rate at which +S+MH is preferred to +S by a nested LRT test. Right column: *ω*_3_ estimates (median, IQR) for various simulation scenarios.

#### Benchmark datasets

We generated an additional 9 null and 18 power simulations (100 replicates each) based on empirical data sets (details shown in Table S1). These scenarios are more representative of biological data because they use alignment sizes, tree topologies, branch lengths, nucleotide substitution biases, and other model parameters based on biological data. We fixed all model parameters except *ω*_3_, *δ*, *ψ*, and *p*_3_. These data recapitulate the patterns found in the simple 4-taxon tree simulations.

1. For null data, +S loses control of FPR as *δ* and/or *ψ* are increased. +S+MH and model averaging maintain FPR control regardless of the values of 2H and 3H rates.
2. For data with EDS but without 2H or 3H, +S has a slight power edge over +S+MH, but model averaging rescues the power because +S has a better goodness of fit.
3. For data with EDS and with 2H and/or 3H, +S has a power edge over +S+MH, and model averaging is only partially able to rescue the power because +S+MH has a much better goodness of fit. This gain in power for +S comes at a cost of significant (often dramatic) upward biases in *ω*_3_ estimates.

Because in real biological data, the presence of selection is the object of inference and it is not expected to be prevalent (e.g., a typical gene is more likely to not be subject to EDS), therefore controlling FP rates should be the prevailing concern. As our simulations demonstrate, unmodeled multi-nucleotide substitutions dramatically inflate the estimates of *ω* rates, and result in significant and often catastrophic FPR.

### A Large-Scale Empirical Screen

We compared the inferences made by the four BUSTED class models on a large-scale empirical dataset (Enard *et al*., 2016) with 9,861 alignments and phylogenetic trees of mammalian species (cf Methods). This collection was originally prepared to assess the influence of viruses on mammalian protein evolution and includes sequences from 24 species.

As with the benchmark datasets, only two out of four model (+S and +S+MH) had the best goodness-of-fit (AICc) measures for most of the alignments (Table 4). BUSTED and +MH were the top model for 97(< 1%) of the alignments which were either very short alignments (<150 codons, e.g., SF3B6 in Table 5) or with minimal divergence (tree length <0.01 substitutions/site). Alignment length and total tree length were not significantly associated (Mann-Whitney U test) with the predilection towards the +S or +S+MH model.

**Table 4.**
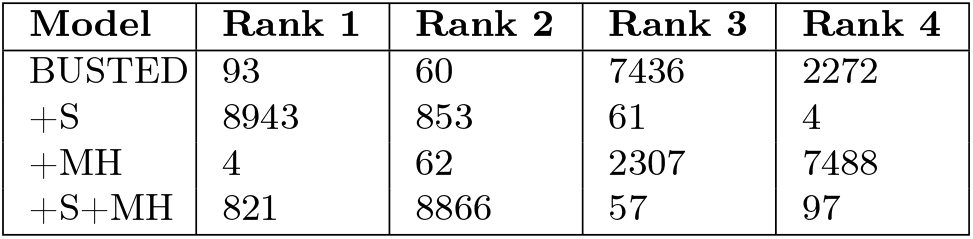
Model goodness-of-fit ranking for the Enard dataset. How each of the four models ranked for each of the 9,861 alignments.

**Table 5.**
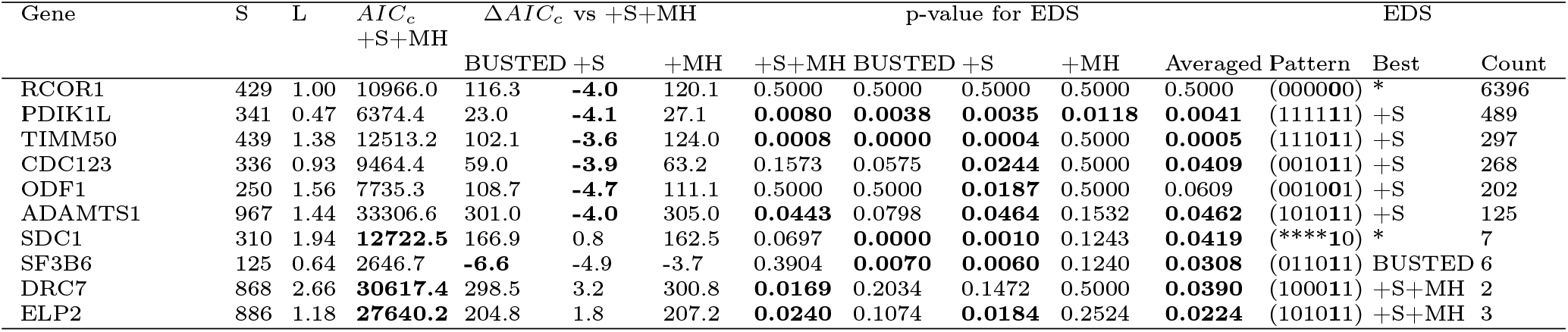
Examples of patterns of agreement and disagreement of different approaches to detecting EDS on the Enard et al dataset, sorted from most to least common. There are 24 sequences in each alignment. **Gene** – gene name, *S* - the number of codons, *L* - total tree length in expected substitutions/nucleotide, measured under the BUSTED+S+MH model. *AIC_c_* S+MH - small sample AIC score for the BUStED+S+MH model (shown in boldface if this model is the best fit for the data, i.e. has the lowest *AIC_c_* score), Δ*AIC_c_* - differences between the *AIC_c_* score for the corresponding model and the BUSTED+S+MH. p-value for ESD: the likelihood ratio test p-value for episodic diversifying selection under the corresponding model (4 digits of precision); shown in boldface if ≤ 0.05. **Averaged** - the model averaged p-value for ESD (bolded if f ≤ 0.05. **Pattern** - a bit vector of whether or not the EDS was detected at *p* ≤ 0.05 with each of the six models: (+S+MH, BUSTED, +S, +MH, Averaged, Best); * denotes a wildcard (0 or 1). **Best**, best fitting model (*AIC_c_*). **Count** - the number of datasets matching this detection pattern.

We next considered whether an alignment was detected subject to EDS, using LRT *p*≤0.05, under different selection criteria: model fixed *a priori*, best fitting-model selected by *AIC_c_*, and the model averaged “p-value” (Fig 4). The simplest model (BUSTED) has the highest raw detection rate of 2805/9861 alignments (28.4%), while the models with MH support have much lower detection rates: 984/9861 (10%) for +S+MH and 826/9861 for +MH (8.4%). Detection of EDS is quite sensitive to which model/approach is being used: there are only 515 alignments, where EDS is detected by all of the models (including the best fitting model), and by model averaging. Requiring a complete model consensus does not strike us as a sensible approach – why, for example, should we give an equal vote to models that do not describe the data well? Indeed, even for the 515 unanimous datasets, the median difference in *AIC_c_* (Δ*AIC_c_*) scores between the best and the worst fitting model was > 200 points, implying that the worst models had much worse fits to the data than the best, and should be discounted. One solution, which has found common use in comparative analysis is to simply pick the best fitting model (such as ModelTest Posada and Buckley (2004)), and call EDS based on it. Here, the best-fitting model detects EDS on 2, 425 datasets. One danger with simply picking the best-fitting model is that in cases when it detects EDS, but the second-best model does not and the second-best model is not dramatically worse fitting, we are discounting a discordant signal from a credible alternative model. The model averaging approach is a simple way to account for this: if two models have similar goodness-of-fit, and one has a low EDS p-value, but the other has a high EDS p-value, averaging the two will result in a non-significant (conservative) call. The “averaging” approach detects EDS on 1,908 datasets. On 524 datasets when the best model detects EDS, but model averaging does not, the median Akaike weight difference *AIC_c_* for the second best fitting model (defined as *w*=*e*^-Δ*AIC_c_*/2^, normalized to sum to 1 over all four models) was 0.18, hence a high p-value from the second-best fitting model is sufficient to push the averaged p-value above 0.05.

**FIG. 4.**
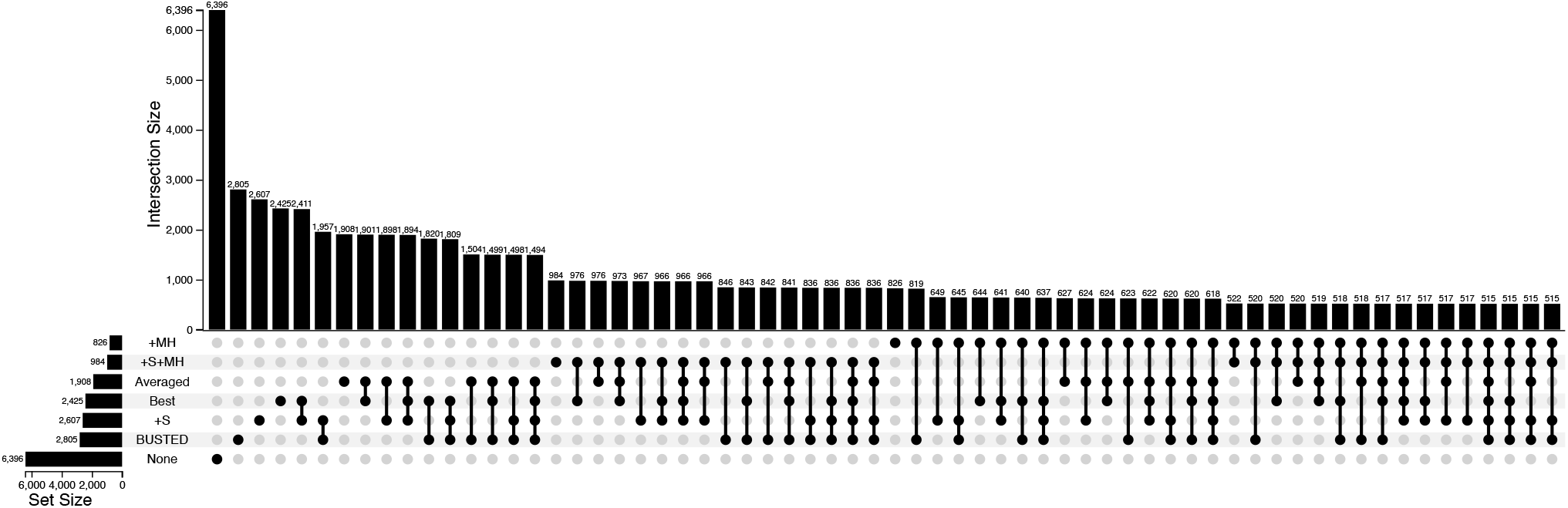
Alignment classification with respect to diversifying selection for the Enard *et al* dataset. The number of alignments (9,861 total) which returned LRT *p*≤0.05 under each of the following scenarios: individual model (BUSTED, +S, +MH, +S+MH), best model selected by *AIC_c_* (Best), model averaged “p-value” (Averaged). The sizes of each of the non-empty intersections of these six combinations are also shown. For example, 515 datasets are found to be subject to EDS under all of the six considered criteria. The category “None” shows all those datasets on which all six approaches failed to achieve significance.

In Table 5, we show examples of patterns for comparative model fit and EDS inference, and discuss them (below) in terms of the selective patterns:

#### No selection detected by any method, pattern (000000)

Nearly two thirds (6396, or 64.9%) of the datasets have no evidence of episodic diversifying selection under any of the six possible detection criteria: (+S+MH, BUSTED, +S, +MH, Averaged, Best). These alignments (e.g., RCOR1), tended to be shorter compared to the alignments with some selection signal (median 427 codons, vs median 599 codons, *p*<10^-16^, Wilcoxson test), have lower overall divergence (median tree length, 1.04 vs 1.27, *p*<10^-16^), and have a smaller fraction of datasets where a model with support for 2H or 3H (odds ratio 0.6, *p*<10^-12^, Fisher exact test).

#### Selection detected by every method (111111)

A total of 515 datasets supported EDS with every detection approach (e.g., PDIK1L), For 489 of those, +S was the best fitting model, and for the remaining 26 – +S+MH was the best fitting model, with longer and more divergent alignments falling into the second bin (+S+MH model), with Wilcoxson p-values of <0.02. Compared to datasets where only some of the methods detected EDS (not consensus), the consensus collection had a larger estimated EDS effect size, approximated by the 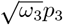 (scaled weight assigned to the positive selection regime), median 0.0133 vs 0.0024 (Wilcoxson *p*=0.001).

#### Selection detected by all but one model

These datasets are “near-consensus” in that all but one of the individual models (e.g., +MH for the (111011) pattern), including the best fitting model and the model-averaged p-value, support EDS (e.g., TIMM50). There are 428 alignments in this bucket, including 401 for which +S is the best fitting model, 25 – +S+MH, and 2 – BUSTED. The most common “outlier” model was +MH (321), followed +S+MH (103), and 2 each for BUSTED and +S.

#### The best model drives EDS selection detection

CDC123 is a prototypical example, where +S is the best model, is the only model that shows evidence of EDS, yet is sufficient for both the Best model and the Averaged model criteria to also indicate EDS. Of the 268 alignments in this group, EDS detection was driven by the +S model for all but two datasets, where the +S+MH model drove detection (DRC7, for example). For all 266 datasets with +S as the best-fitting model, +S+MH was the second-best model, receiving a median Akaike weight of only 0.046, i.e. making it irrelevant for model averaged p-value calculations.

#### +S and +S+MH models both detect EDS

For genes like ADAMTS1and ELP2, +S and +S+MH are the two credible models, that both detect EDS, together with the Best and Averaged approaches. There are 125 of such datasets for which +S is the best-fitting model, and 3 with +S+MH as the best-fitting model.

#### The best model drives EDS selection detection, but model averaging disagrees

The first class of datasets where important disagreement occurs, are those where EDS is detected with the best fitting model but not detected with the model averaged approach. ODF1 is an example: the best-fitting model (+S) supports EDS with *p*=0.0187, but the second-best model (+S+MH), finds no evidence EDS (*p*=0.5). Model-averaging takes both of those indications into account and arrives at a non-significant p-value of 0.06. There are 524 datasets in this bucket, and for all but 9 of those, +S is the best model, and +S+MH is second best and plays the role of spoiler. Median MLEs for MH rates were higher in the datasets than where +S and +S+MH disagreed (only +S supports EDS), compared to where they agreed (both models support EDS): 0.03 vs 0.0, *p*<10^-10^ for δ; 0.03 vs 0.05, *p*<10^-10^ for ψ). The models also had significantly different (*p*<10^-10^) estimates for *ω_3_* with median differences 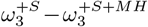 of 24.7 (EDS only for +S) vs 0.01 (EDS in both). These patterns are consistent with false positive EDS detection by the +S model as seen on simulated data.

#### Model averaging finds EDS, but the best-fitting model disagrees

There are only 7 datasets (e.g., SDC1), in this counter-intuitive class of an important disagreement. For these types of datasets, the best fitting model has a borderline significant p-value, the second best-fitting model has a highly significant p-value (and is very close in terms of *AIC_c_*), and the model averaging approach arrives at a significant p-value.

#### The effect of EDS detection strategies under muptiple testing correction

Depending on the specific type of analysis, large-scale screens of genes for a specific feature (e.g. EDS) often require some form of multiple-testing correction. Such correction can be directly applied to p-values from of the EDS detection methods we described above. For example, Table 6 shows the rates of EDS detection for all 6 methods considered here using raw p-values (*p*≤0.05) and also false-discovery rate corrected q-values (Benjamini-Hochberg FDR, *q*≤0.2). For models without MH (BUSTED, +S), the effect of FDR correction is effectively nil, at least for these thresholds, whereas for models with MH (+MH, +MH+S), performing FDR dramatically reduces already low detection rates by about 3-fold. Encouragingly, the model-averaged approach is much less affected by the FDR correction, with the detection rate dropping from 19.3% to 15.5%.

**Table 6.** EDS detection rates with and without multiple testing correction.

| Method | $p \leq 0.05$ | $q \leq 0.20$ |
| --- | --- | --- |
| BUSTED | 0.284 | 0.2985 |
| +S | 0.264 | 0.272 |
| +MH | 0.0838 | 0.0271 |
| +S+MH | 0.0998 | 0.0348 |
| Best model | 0.246 | 0.244 |
| Model averaged | 0.193 | 0.155 |

## Discussion

Evolutionary substitution models that are practically useful and computable must make many simplifying assumptions about biological processes. Many, if not most, of these assumptions are not justifiable on biological grounds. However, certain classes of inference problems appear to be quite robust to even severe model misspecifications. Examples include phylogenetic inference (Abadi *et al*., 2019), and relative evolutionary rate estimates for individual sites (Spielman and Kosakovsky Pond, 2018). Other inference problems, including selection detection, seem to be highly sensitive to modeling assumptions (Kosakovsky Pond *et al*., 2011; Venkat *et al*., 2018). Such sensitivity is not surprising for methods that are tuned to extract statistical signal from a small subset of branches and sites in a sequence alignment. In extreme cases, a single substitution event is sufficient to power selection detection, as seen in the HIV-1 *v*if example in this paper and for human lineage selection detection in Venkat *et al*. (2018).

Statistical tests which compare the *ω* ratio of non-synonymous and synonymous substitution rates to 1 and interpret significant differences as evidence of non-neutral evolution are susceptible to confounding processes which bias *ω* estimates. We have previously demonstrated that *ω* estimates are strongly biased (and resulting in high Type 1 and 2 error rates) when the distribution used to model *ω* variation across branches and sites is too restrictive (Kosakovsky Pond *et al*., 2011), and when synonymous substitution rates are assumed to be constant across sites in the alignment (Wisotsky *et al*., 2020). Furthermore, these confounding processes are not rare, but instead are very likely present in biological data. Because “the scientist must be alert to what is importantly wrong” (Box, 1976), and these models are clearly wrong in important ways, as they misinterpret widespread confounding evolutionary processes as evidence of selection, continued use of such models is unsound.

Using simulations and empirical data, our study corroborates the conclusions of multiple previous reports (e.g., Venkat *et al*. (2018); Whelan and Goldman (2004); Wisotsky *et al*. (2020)) on the impact that MH has on evolutionary process inference and characterization. Not accounting for instantaneous multi-nucleotide substitutions or “hits” (MH) when estimating evolutionary process parameters, especially related to natural selection, leads to serious statistical misbehavior of widely used tests, and biases rate estimates. Estimates of *ω* become inflated with standard codon substitution models when they are used to analyze data with MH, and progressively more so as the degree of MH is increased. This bias, in turn, produces uncontrolled rates of false positives for positive selection on simulated data for MH parameter values that appear realistic. These findings add to the already significant body of literature in this space. This is the first study, however, to demonstrate how to model MH and synonymous site-to-site rate variation (SRV) jointly, and that MH remains an important factor even if SRV is accounted for.

In a large-scale empirical analysis of mammalian genes (Enard *et al*., 2016) we found that 10% of alignments are best fit with models supporting MH, and that roughly 80% of positively selected genes are robustly detected even when accounting for MH using a model-averaging procedure. Consequently, confounding due to MH can be viewed as a “second-order” effect, compared, for example, to the inclusion of synonymous site-to-site rate variation (SRV), which impacts the vast majority of genes. However, we argue that even second-order effects are sufficiently important to be considered in routine analyses of selection. Our practical recommendation, supported by simulated data and empirical analyses, is to fit multiple flavors of selection models followed up by model-averaged selection detection to obtain a good tradeoff between power and false positive rate control. We also developed a series of visual tools to assist researchers in interpreting selection analysis results, exploring which branches and sites in the alignment provide support for various evolutionary processes (selection and/or MH), and understanding how much a positive selection result is influenced by information from a small number of sites.

There are other forms of potential model violation that we did not address here, which could also bias tests of EDS. For example, MNMs are known to be enriched for transversions, which tend to be nonsynonymous, and failure to incorporate this bias has been shown previously to inflate false positive rates of EDS tests Venkat *et al*. (2018). Other potentially important forms of complexity are heterogeneity in preference among amino acids and heterogeneity among pairs, e.g., Dunn *et al*. (2019), as well as among-site differences in these factors (e.g., Bloom (2014)). These approaches broadly fall into the category of models where the *ω* ratio is a function of the aminoacids being exchanged (Delport *et al*., 2010b). The key issue for implementing such models is selecting a biologically or empirically justified parameterization, which remains an open problem because it is relatively trivial to improve the goodness-of-fit over the baseline model (Delport *et al*., 2010a), and such improvement should not be viewed as *prima facie* evidence of biological significance. Further research is necessary to incorporate these and other kinds of complexity into tractable Markov models and/or evaluate their effects on EDS analyses.

A plethora of studies (Cohen *et al*., 2021; Freitas and Nery, 2022; Hensley *et al*., 2021; Lucaci *et al*., 2021; MacLean *et al*., 2021; Steward *et al*., 2022; Venkat *et al*., 2018) suggest that MH occurs broadly over diverse taxonomic groups. We expect that future research with an interdisciplinary design combining computational and experimentally-informed approaches may shed light on the application of our method(s) and the patterns and processes underlying the contribution of MH to gene evolution. Creative investigation may help discover additional mechanisms and interpretations of the biological underpinnings of the mutational spectrum as it applies to rare mutations in natural populations. Additionally, we see a strong tailwind in this field as technological improvements for functional studies designed with the precise manipulation of DNA (Wang *et al*., 2020) including CRISPR-Cas9, and detection of MH polymorphisms (J. Huang *et al*., 2014) continue to emerge, draw interest, and be fine-tuned. Downstream innovations and technological design are critical in the detection of natural selection, where models such as ours can be of particular interest to researchers interested in gene-drug target design for particular fitness effects. Additionally, our work supports an emerging body of information on the underlying trends, biological mechanisms, and genetic signaling pathways under selective pressure. These results can feed directly into a number of *post hoc* analyses to qualify or quantify an exploratory genetic profile and evolutionary history across lineages.

## Methods

### Statistical Methodology

We adapted two existing models: the BUSTED model, a test of episodic diversifying selection, by (Murrell *et al*., 2015), and the +S model by (Wisotsky *et al*., 2020), which was created as a modification of the BUSTED model, to account for the presence of synonymous rate variation (SRV). The +S+MH model is a straightforward extension of +S which allows it to account for instantaneous multiple nucleotide changes occurring within a codon (MH) and SRV, while the BUSTED+MH model is an extension of the BUSTED model where SRV is not modeled (Table 1). In this framework, the nucleotide substitution process is described using the standard discrete-state continuous-time Markov process approach of (Muse and Gaut, 1994), with entries in the instantaneous rate matrix (*Q*) corresponding to substitutions between sense codons *i* and *j* and defined as follows:

Here, *θ_ij_*(=*θ_ji_*) denote nucleotide substitution bias parameters. For example, *θ_ACT,AGT_*=*θ_CG_* and because we incorporate the standard nucleotide general time-reversible (GTR) (Tavare, 1986) model there are five identifiable *θ_ij_* parameters: *θ_AC_*, *θ_AT_*, *θ_CG_*, *θ_CT_*, and *θ_GT_* with *θ_AG_*=1. The position-specific equilibrium frequency of the target nucleotide of a substitution is 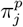; for example, it is 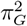 for the second-position change associated with *q_ACT,AGT_*. The 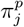 and the stationary frequencies of codons under this model are estimated using the CF3×4 procedure (Pond *et al*., 2010), adding nine parameters to the model. The ratio of nonsynonymous to synonymous substitution rates for site *s* along branch *b* is *ω^bs^*, and this ratio is modeled using a 3-bin general discrete distribution (GDD) with five estimated hyperparameters: 0≤*ω*_1_≤*ω*_2_≤1≤*ω*_3_, *p*_1_ = *P*(*ω^bs^* = *ω*_1_), and *p*_2_ = *P*(*ω^bs^* = *ω*_2_) (Table 7). The procedure for efficient computation of the phylogenetic likelihood function for these models was described in Kosakovsky Pond *et al*. (2011).

**Table 7.** BUSTED+S+MH rate matrix definition.

| Type | Expression for $Q_{ij}$ |
| --- | --- |
| 1 step synonymous change | $\alpha^s \theta_{ij} \pi_j^p$ |
| 1 step nonsynonymous change | $\alpha^s \omega^{bs} \theta_{ij} \pi_j^p$ |
| 2 step synonymous change | $\delta \alpha^s \prod_{n=1}^2 \theta_{ij}^n \pi_j^n$ |
| 2 step nonsynonymous change | $\delta \alpha^s \omega^{bs} \prod_{n=1}^2 \theta_{ij}^n \pi_j^n$ |
| 3 step synonymous change | $\psi \alpha^s \prod_{n=1}^3 \theta_{ij}^n \pi_j^n$ |
| 3 step nonsynonymous change | $\psi \alpha^s \omega^{bs} \prod_{n=1}^3 \theta_{ij}^n \pi_j^n$ |

The quantity *α^s^* is a site-specific synonymous substitution rate (no branch-to-branch variation is modeled) drawn from a separate 3-bin GDD. The mean of this distribution is constrained to one to maintain statistical identifiability, resulting in four estimated hyperparameters: 0≤*cα*_1_≤*α*_2_=*c*≤*cα*_3_, *f*_1_=*P*(*α^s^*=*α*_1_), and *f*_2_=*P*(*α^s^*=*α*_2_), with *c* chosen to ensure that E[*α_s_*]=1.

The key parameters are global relative rates of multiple hit substitutions: *δ* is the rate for two substitutions relative to the one substitution synonymous rate (baseline), *ψ* is the relative rate for non-synonymous three substitutions. All parameters, except *π*, including branch lengths, are fitted using a directly optimized phylogenetic likelihood in HyPhy.

Typical implementations, including ours, allow the number of *α* and *ω* rate categories to be separately adjusted by the user, for example, to minimize cAIC or to optimize some other measure of model fit. The default setting of three categories generally provides a good balance between fit and performance when using this GDD approach for modeling. Our implementation of +S+MH, and BUSTED+MH will warn the user if there is evidence of model overfitting, such as the appearance of rate categories with very similar estimated rate values or very low frequencies.

A notable limiting assumption of the model, made for computational tractability, is that MH substitutions spanning codon boundaries, e.g. a substitution in the third position in a codon coupled with a substitution in the first position of the subsequent codon, are not accounted for. Doing so in a systematic fashion breaks the assumption of site independence, making computation largely intractable. One approximate solution is a mean-field type approximation (Whelan and Goldman, 2004), but even this approach is computationally quite expensive. Not accounting for “codon-spanning” multiple hits is a conservative approach, since it will *miss* a subset of MH substitutions, treating them instead as several single nucleotide substitutions occurring independently.

The +S+MH procedure for identifying positive selection is the likelihood ratio test comparing the full model described above to the constrained model formed when *ω*_3_ is set equal to 1 (i.e., no positively selected sites). Critical values of the test are derived from a 50:50 mixture distribution of 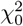 and 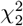 (Murrell *et al*., 2015; Wisotsky *et al*., 2020). Both +S and +S+MH analyses in the current work use the same 50:50 mixture test statistic. +S+MH reduces to +S by setting the MH rates to 0. The method is implemented as a part of HyPhy (version 2.5.42 or later) (Kosakovsky Pond *et al*., 2020).

### Empirical Data and Alignments

The (Enard *et al*., 2016) data collection includes 9,861 orthologous coding sequence alignments of 24 mammalian species and is available at https://datadryad.org/stash/dataset/doi:10.5061/dryad.fs756. Phylogenetic trees were inferred for each alignment using RAxML (Kozlov *et al*., 2019).

### Synthetic data

Simulated data sets can be downloaded from https://data.hyphy.org/web/busteds-mh/.

Additional information is present in the `README.md` file, including details of how to generate alignments under the +S and +S+MH models.

### Implementation

All analyses were performed in HyPhy version 2.5.42 or later. The BUSTED+MH and +S+MH models are implemented as part of the standard HyPhy library. You can run this option using the “-multiple-hits” option from the command line with either “Double” to consider DH substitutions or “Double+Triple” to consider DH and TH substitutions. The HyPhy Batch Language (HBL) implementation is located in a dedicated GitHub repository at https://github.com/veg/hyphy

### Site-level support

In order to identify which individual sites show preference for MH models, we use evidence ratios (ER), defined as the ratio of site likelihoods under two models being compared. We previously showed that ERs are useful for identifying the sites driving support for one model over another, and they incur trivial additional overhead to compute once model fits have been performed.

### Empirical Bayes support

We can estimate statistical support for selection (*ω*_3_>1) or multiple hit substitutions (*δ*>0 or *ψ*>0) at a particular site (*s*) and branch (*b*), using a straightforward empirical Bayes calculation. For example, 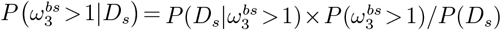, where *P*(*D_s_*) is the standard phylogenetic likelihood of *D_s_* (summed over all *ω* combinations), 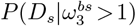) is the phylogenetic likelihood at site *s*, computed by setting the distribution of *ω* at branch *b* to assign all weight to *ω*_3_ > 1, and 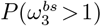 is the mixture weight estimated from the entire alignment (MLE for the corresponding hyperparameter). The corresponding empirical Bayes factor (EBF) is 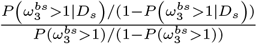. As discussed in Murrell *et al*. (2012), these empirical estimates are quite noisy and should only be used for exploratory purposes, e.g., to look for “hot-spots” in a tree (cf Figure 1).

### Hypothesis testing

Nested models are compared using likelihood ratio tests with asymptotic distribution used to assess significance. A conservative 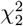 asymptotic distribution is used to compare the fit of +S and +S+MH (null hypothesis: *δ*=*ψ*=0).

### Computational complexity

Treating BUSTED as a baseline, we expect the +S model to require about a relative ×*L* (L = number of synonymous rate classes) more time per likelihood calculation and longer convergence time due to an extra random effects distribution.

Because +MH models have dense rate matrices, there is a computational cost incurred for computing transition matrices since optimizations available for standard (sparse) matrices no longer apply. +S+MH models are expected to be the slowest, but have the same order of complexity as +*S*. On 24-sequence alignments from Enard *et al*. (2016), we observed the following performance for each of the four models (Table 8).

**Table 8.**
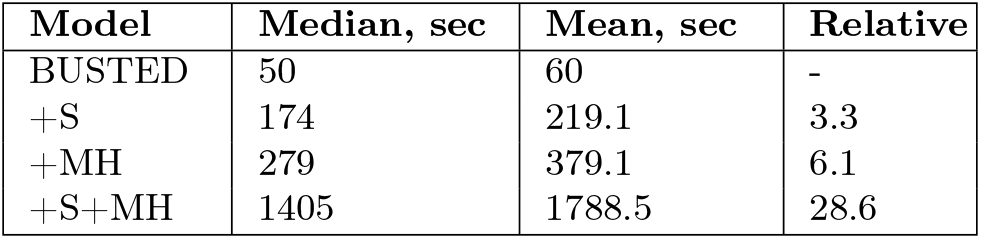
Model run-times for the Enard dataset. Run times on 4 cores on an AMD EPYC 7702 CPU compute node are shown. `Relative`: median of the relative run times on the same dataset compared to BUSTED.

### BUSTED ModelTesting

We recommend our BUSTED model testing and averaging procedure (see main text) in order to select the best fitting model, to interpret the results of natural selection acting on your gene of interest. Our goal is to understand which underlying model and its parameters are able to detect the areas of the dataset which drive the greatest degree of evolutionary signals. We screen the dataset for episodic diversifying selection acting on the whole gene while accounting for SRV across the alignment, and MH substitutions.

Analysis is conducted as a series of experiments in the BUSTED framework of selection analysis with our methods under analysis in the hierarchical structure described in Table 1 and includes BUSTED, +S, BUSTED+MH, and +S+MH.

We implement a Snakemake (Mölder *et al*., 2021) version of our model testing procedure, available at https://github.com/veg/BUSTED_ModelTest. This application takes the same input as a normal BUSTED analysis, a multiple sequence alignment and inferred phylogenetic, and returns JavaScript Object Notation (JSON) files (one for each model described above).

We recommend performing model averaging to determine whether or not an alignment is subject to episodic diversifying selection. Alternative approaches could include selecting the best fitting model, or model consensus, however, as shown by our simulations, these approaches are less statistically efficient (lower power and/or higher rate of false positives).

## Acknowledgments

We thank members of the HyPhy and Datamonkey teams for their contribution to this work. This research was supported in part by grants GM144468 (NIH/NIGMS), AI140970 (NIH/NIAID), AI134384 (NIH/NIAID), and NIH NIGMS R35GM142677 (Enard).

