## Supplementary material for "Evolutionary shortcuts via multi-nucleotide substitutions and their impact on natural selection analyses": NA

· doi:10.1093/molbev/mst012 MBE

### Supplementary Material

· doi:10.1093/molbev/mst012 MBE

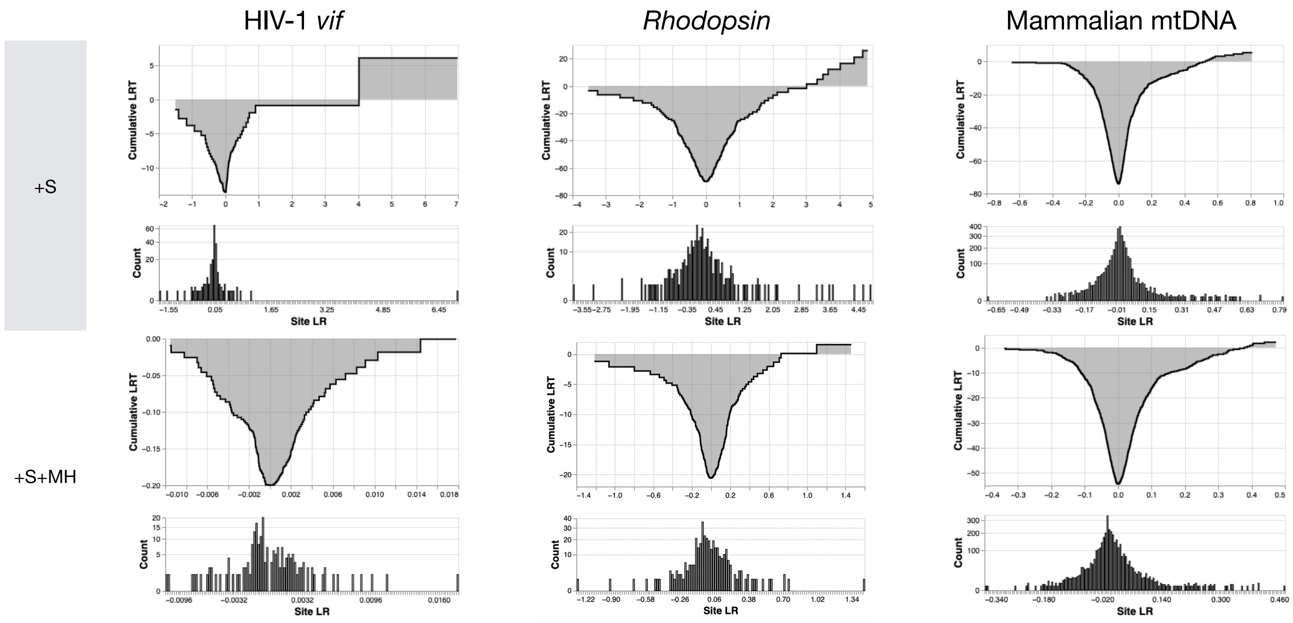


**FIG. S1. Site-level support for Episodic Diversifying Selection in three benchmark alignments**. Each dataset / model panel includes two views of the same data: the top plot is the cumulative value of the likelihood ratio test statistic (LRT) for the EDS test over sites, where site-level LRT are sorted from smallest to largest; the bottom plot is the histogram of site-level LRTs.

| Scenario | $\omega_3(p_3)$ | | | $\delta$ | | $\psi$ | | Detection (# if $AIC_c$ is better) | | | +S+MH pref |
|---|---|---|---|---|---|---|---|---|---|---|---|
| | Truth | +S | +S+MH | Truth | +S+MH | Truth | +S+MH | +S | +S+MH | Averaged | by LRT |
| **Null simulations (no positive selection, MH present)** | | | | | | | | | | | |
| adh/N1 | **1.0**(2.38%) | 1.00 − 2.42 (1.6%) | 1.00 − 2.63 (1%) | 0.003 | 0.00 − 0.04 | 0.0 | 0.00 − 0.00 | 0.03 (97) | 0.01 (3) | 0.01 | 0 |
| adh/N2 | **1.0**(2.38%) | 1.00 − 3.60 (1.7%) | 1.00 − 1.93 (0.98%) | 0.1 | 0.00 − 0.12 | 0.0 | 0.00 − 0.07 | 0.1 (80) | 0 (20) | 0.04 | 0.14 |
| adh/N3 | **1.0**(2.38%) | 1.30 − 5.21 (2.3%) | 1.00 − 2.26 (1.2%) | 0.25 | 0.10 − 0.27 | 0.0 | 0.00 − 0.08 | 0.26 (47) | 0.03 (53) | 0.05 | 0.39 |
| adh/N4 | **1.0**(2.38%) | 2.07 − 7.98 (2.3%) | 1.00 − 2.69 (1.3%) | 0.5 | 0.33 − 0.56 | 0.0 | 0.00 − 0.06 | 0.61 (6) | 0.02 (94) | 0.02 | 0.9 |
| adh/N5 | **1.0**(2.38%) | 3.05 − 19.29 (2.1%) | 1.00 − 2.32 (1.1%) | 0.75 | 0.57 − 0.79 | 0.0 | 0.00 − 0.12 | 0.87 (0) | 0.03 (100) | 0.03 | 1 |
| Hepatitis D Ag/N1 | **1.0**(1.71%) | 1.19 − 10.93 (13%) | 1.00 − 1.04 (6.9%) | 0.14 | 0.08 − 0.15 | 0.0 | 0.00 − 0.00 | 0.34 (32) | 0.01 (68) | 0.06 | 0.54 |
| HIV *vif*/N1 | **1.0**(1.00%) | 1.07 − 50.21 (14%) | 1.00 − 1.90 (7.2%) | 0.004 | 0.00 − 0.00 | 0.16 | 0.08 − 0.22 | 0.32 (32) | 0.01 (68) | 0.01 | 0.63 |
| Rhodopsin/N1 | **1.0**(0.37%) | 3.37 − 14.11 (1.1%) | 1.00 − 1.69 (0.6%) | 0.35 | 0.27 − 0.37 | 0.52 | 0.27 − 0.64 | 0.9 (1) | 0.02 (100) | 0.03 | 0.99 |
| Strep. PTS/N1 | **1.0**(1.56%) | 1.48 − 12.41 (1.6%) | 1.00 − 5.59 (0.93%) | 0.31 | 0.15 − 0.40 | 1.1 | 0.70 − 1.46 | 0.31 (1) | 0.03 (99) | 0.03 | 0.99 |
| **Power simulations (positive selection, MH absent)** | | | | | | | | | | | |
| adh/P1 | 4.14(2.50%) | 3.21 − 4.79 (2.9%) | 3.30 − 4.91 (2.8%) | 0.0 | 0.00 − 0.00 | 0.0 | 0.00 − 0.00 | 0.91 (99) | 0.85 (1) | 0.92 | 0.01 |
| $\beta$-globin/P1 | 8.925(3.70%) | 5.53 − 10.96 (5%) | 5.73 − 9.67 (4.8%) | 0.0 | 0.00 − 0.00 | 0.0 | 0.00 − 0.03 | 1 (91) | 0.98 (9) | 1 | 0.05 |
| HIV *vif*/P1 | 2103(0.05%) | 1.09 − 3142.84 (9.7%) | 1.00 − 2.54 (6.2%) | 0.0 | 0.00 − 0.02 | 0.0 | 0.00 − 0.11 | 0.56 (80) | 0.1 (20) | 0.53 | 0.14 |
| Mam. mtDNA/P1 | 1.434(1.33%) | 1.29 − 1.63 (1.3%) | 1.21 − 1.49 (1.4%) | 0.0 | 0.00 − 0.03 | 0.0 | 0.00 − 0.05 | 0.65 (90) | 0.5 (10) | 0.64 | 0.06 |
| Rhodopsin/P1 | 6.376(1.31%) | 5.13 − 7.38 (1.4%) | 5.16 − 7.50 (1.4%) | 0.0 | 0.00 − 0.00 | 0.0 | 0.00 − 0.10 | 1 (99) | 0.98 (2) | 0.99 | 0.01 |
| **Power simulations (positive selection, MH present)** | | | | | | | | | | | |
| adh/P2 | 4.05(2.38%) | 3.23 − 5.40 (2.6%) | 3.04 − 5.05 (2.5%) | 0.003 | 0.00 − 0.03 | 0.0 | 0.00 − 0.06 | 0.9 (94) | 0.75 (6) | 0.86 | 0.02 |
| $\beta$-globin/P2 | 2.834(6.08%) | 2.71 − 6.40 (7%) | 1.88 − 4.54 (7.4%) | 0.24 | 0.01 − 0.28 | 0.19 | 0.00 − 0.14 | 0.88 (56) | 0.4 (44) | 0.55 | 0.33 |
| Hepatitis D Ag/P1 | 11.3(1.71%) | 8.93 − 19.65 (3.7%) | 2.40 − 9.86 (9.4%) | 0.14 | 0.07 − 0.17 | 0.0 | 0.00 − 0.07 | 1 (32) | 0.58 (68) | 0.65 | 0.57 |
| HIV *vif*/P2 | 1.226(1.00%) | 1.64 − 319.54 (9.4%) | 1.00 − 3.12 (6.2%) | 0.004 | 0.00 − 0.00 | 0.16 | 0.08 − 0.23 | 0.46 (35) | 0.01 (65) | 0.04 | 0.52 |
| Rhodopsin/P2 | 5.453(0.37%) | 5.61 − 15.94 (0.96%) | 1.00 − 6.27 (0.73%) | 0.35 | 0.26 − 0.35 | 0.52 | 0.25 − 0.81 | 1 (1) | 0.24 (100) | 0.25 | 0.99 |
| Rhodopsin/P3 | 5.453(0.37%) | 4.14 − 9.24 (1.1%) | 1.00 − 5.47 (0.61%) | 0.35 | 0.26 − 0.37 | **0.0** | 0.00 − 0.18 | 0.96 (0) | 0.22 (100) | 0.22 | 0.97 |
| Rhodopsin/P4 | 5.453(0.37%) | 3.59 − 8.65 (0.72%) | 2.23 − 7.11 (0.71%) | **0.10** | 0.03 − 0.13 | **0.0** | 0.00 − 0.08 | 0.87 (70) | 0.32 (30) | 0.55 | 0.18 |
| Rhodopsin/P5 | 5.453(0.37%) | 3.15 − 8.12 (1.1%) | 1.37 − 6.64 (0.83%) | **0.20** | 0.14 − 0.24 | **0.0** | 0.00 − 0.09 | 0.85 (21) | 0.27 (79) | 0.37 | 0.7 |
| Rhodopsin/P6 | 5.453(0.37%) | 4.47 − 20.37 (0.6%) | 1.62 − 6.06 (0.71%) | **0.00** | 0.00 − 0.01 | 0.52 | 0.35 − 0.60 | 0.96 (26) | 0.28 (74) | 0.39 | 0.62 |
| Rhodopsin/P7 | 5.453(**2.1**%) | 7.21 − 11.00 (2.2%) | 4.65 − 6.46 (2.2%) | 0.35 | 0.28 − 0.38 | **0.0** | 0.00 − 0.12 | 1 (3) | 0.98 (97) | 0.99 | 0.97 |
| Rhodopsin/P8 | 5.453(**4.2**%) | 7.51 − 9.64 (4.1%) | 4.78 − 6.15 (4.4%) | 0.35 | 0.26 − 0.36 | **0.0** | 0.00 − 0.06 | 1 (3) | 0.99 (97) | 1 | 0.97 |
| Strep. PTS/P1 | 9.489(1.56%) | 17.10 − 71.54 (1.4%) | 7.55 − 12.88 (1.6%) | 0.31 | 0.15 − 0.43 | 1.1 | 0.65 − 1.45 | 0.99 (0) | 0.97 (100) | 0.97 | 1 |
| SARS-CoV-2 S/P1 | 5.990(20.12%) | 4.64 − 7.63 (27%) | 4.86 − 8.94 (22%) | 0.012 | 0.00 − 0.00 | 0.0 | 0.00 − 0.00 | 0.99 (100) | 0.97 (0) | 0.99 | 0 |

**Table S1.** BUSTED test performance on synthetic data, under model fits from benchmark datasets to parametrize various simulation scenarios simulations (100 replicates each). `Truth` – values used for data generation; parameters changed from their MLE values from the corresponding empirical dataset are shown in **boldface**. For model rate estimates, interquartile range is shown. For proportion estimates, mean value is shown. `Detection` columns shows the fraction of replicates where the LRT for episodic diversifying selection yields $p \leq 0.05$; and the value in parentheses – the number of replicates where the corresponding model was preferred by $AIC_c$. `Detection / Averaged` – the fraction of replicates where model-averaged LRT p-value was $\leq 0.05$. The last column shows the fraction of replicates for which the +S+MH model was preferred to the +S model, using the $\chi^2_2$ based LRT $p \leq 0.05$.

· doi:10.1093/molbev/mst012 MBE

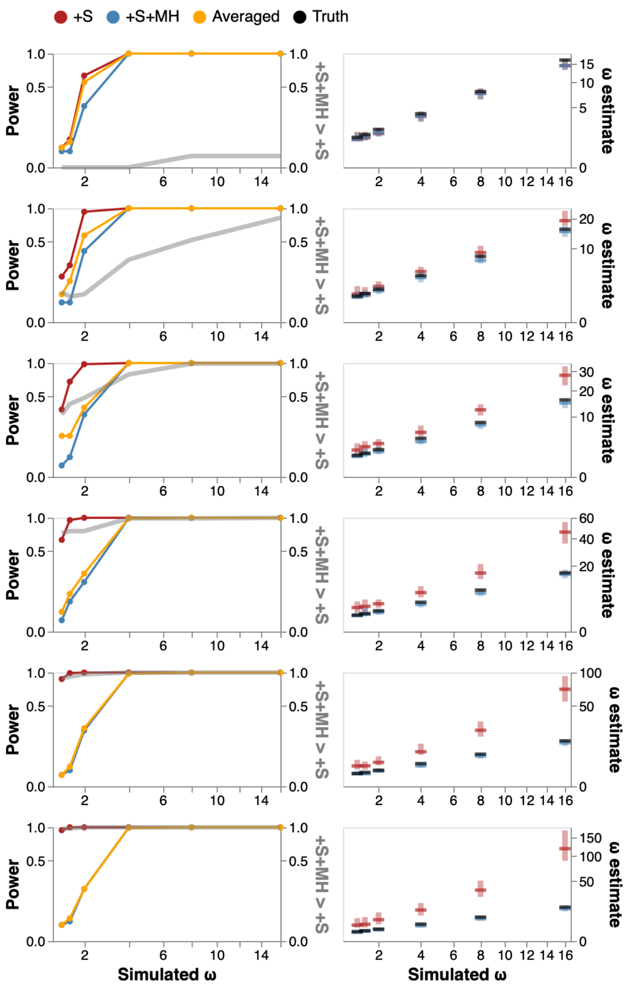


**FIG. S2. Model performance on data simulated with EDS (25% selected fraction).** Left column : detection rate for EDS (at $p \leq 0.05$) as a function of rate $\omega_3$ (effect size) and $\delta$ (confounding parameter), and the rate at which +S+MH is preferred to +S by a nested LRT test. Right column: $\omega_3$ estimates (median, IQR) for various simulation scenarios.
